# Spherical Spindle Shape Promotes Perpendicular Cortical Orientation by Preventing Isometric Cortical Pulling on both Spindle Poles during *C. elegans* Female Meiosis

**DOI:** 10.1101/596510

**Authors:** Elizabeth Vargas, Karen P. McNally, Daniel B. Cortes, Michelle T. Panzica, Amy Shaub-Maddox, Francis J. McNally

## Abstract

Meiotic spindles are positioned perpendicular to the oocyte cortex to facilitate segregation of chromosomes into a large egg and a tiny polar body. In *C. elegans*, spindles are initially ellipsoid and parallel to the cortex before shortening to a spherical shape and rotating to the perpendicular orientation by dynein-driven cortical pulling. The mechanistic connection between spindle shape and rotation has remained elusive. Here we used mutants of the microtubule-severing protein katanin to manipulate spindle shape without eliminating cortical pulling. In a katanin mutant, spindles remained ellipsoid, had pointed poles and became trapped in either a diagonal or a parallel orientation. Results indicated that astral microtubules emanating from both spindle poles initially engage in cortical pulling until microtubules emanating from one pole detach from the cortex allowing pivoting of the spindle. The lower viscous drag experienced by spherical spindles prevented recapture of the cortex by astral microtubules emanating from the detached pole. In addition, maximizing contact between pole dynein and cortical dynein stabilizes round poles in a perpendicular orientation. Spherical spindle shape can thus promote perpendicular orientation by two distinct mechanisms.

## Introduction

The vast majority of eukaryotes are diploid and reproduce sexually using the conserved process of meiosis. Oocytes of most animals undergo a highly asymmetric meiosis in which only one of four sets of maternal chromosomes is inherited by a zygote. Female meiotic spindles are positioned with one pole closely apposed to the oocyte cortex during anaphase. This perpendicular orientation of the meiotic spindle facilitates asymmetric cell divisions that deposit half the homologous chromosomes in a first polar body and half the remaining sister chromatids in a second polar body (Fabritius et al., 2011a).

In *C. elegans*, the precursor of the meiotic spindle, the germinal vesicle, migrates from the center of the oocyte to the cortex through a mechanism that requires kinesin-1 heavy chain (UNC-116) and its binding partner KCA-1 (Yang et al., 2005; McNally et al., 2010). At nuclear envelope breakdown, the spindle assembles near the cortex, then migrates an average of 2 μm towards the cortex and adopts a roughly parallel orientation to it. The spindle maintains this parallel orientation and an ellipsoid spindle shape (axial ratio 1.3) with a steady state length of 8 μm, for an average of 7 minutes. Then, in a stereotypical series of anaphase promoting complex (APC)-dependent events (Yang et al., 2003; Yang et al., 2005), the spindle shortens in the pole to pole axis, achieving an axial ratio 1.0, then rotates to a perpendicular orientation with one pole moving closer to the cortex (Crowder et al., 2015). Anaphase A chromosome separation initiates during or just after rotation as the spindle continues to shorten to an axial ratio of 0.8. Upon completion of anaphase A, the perpendicular spindle elongates and transforms into a cylindrical shape as anaphase B progresses. Finally, the half spindle proximal to the cortex is pushed through the cytokinetic ring (Dorn et al. 2010) or the cytokinetic ring ingresses to the spindle midpoint (Fabritius et al., 2011b) and the chromosomes at the cortex are partitioned into a polar body.

In embryos depleted of either kinesin-1 or KCA-1, nuclear migration and early translocation of the spindle to the cortex fail, resulting in metaphase spindles that are, on average, 7 μm from the cortex (McNally et al., 2010; Yang et al., 2005). However, upon APC activation, as the spindle begins to shorten, it translocates to the cortex. This late, compensatory translocation in kinesin-depleted embryos appears to occur by the same mechanism as spindle rotation in control embryos because both movements require the anaphase promoting complex (Yang et al., 2005), CDK1 inactivation (Ellefson and McNally, 2011) and cytoplasmic dynein (van der Voet et al., 2009; Ellefson and McNally, 2009). Initiation of dynein-dependent spindle rotation correlates with spherical spindle shape even when spindle shape is manipulated by changing ploidy (Crowder et al., 2015) but the mechanistic reason for the correlation between shape and orientation has remained obscure.

Well characterized dynein-dependent spindle positioning mechanisms involve astral microtubules with minus ends attached to a centrosome or spindle pole body and plus ends that interact with cytoplasmic dynein anchored at the cell cortex (McNally, 2013; Kotak, 2019). The mechanism of dynein-dependent meiotic spindle positioning is less understood because, while *C. elegans* meiotic spindles have astral microtubules and cortical dynein (Crowder et al., 2015), they have no centrosomes (Albertson and Thomson, 1993). Therefore it is unclear how the minus ends of astral microtubules would be anchored in an acentrosomal spindle pole to allow pulling by cortical dynein. *C. elegans* meiotic embryos also have cortical microtubules and these microtubules are inferred to have minus ends anchored at the cortex and plus ends oriented inward because they allow kinesin-1 dependent packing of yolk granules into the embryo interior (McNally et al., 2010; Kimura et al., 2017). Plus ends of the cortical microtubules might interact with dynein that accumulates on spindle poles during rotation (Ellefson and McNally, 2011). However, it is unclear how dynein could be anchored at an acentrosomal spindle pole in a manner that would allow pulling on the plus ends of cortical microtubules. Recently, Tan et al. (2018) demonstrated that when a threshold number of dynein, dynactin, and bicD complexes accumulate at a microtubule minus end, they acquire the ability to capture a second microtubule and move the minus end of the first microtubule toward the minus end of the second microtubule. This mechanism could explain how spindle pole dynein could pull on the plus ends of cortical microtubules or how the minus ends of astral microtubules would be mechanically attached to the spindle when pulled by cortical dynein. Meiotic spindle rotation requires ASPM-1, LIN-5, dynein and dynactin (van der Voet et al., 2010; Crowder et al., 2015) and it is possible that ASPM-1/LIN-5 substitutes for BicD during spindle rotation to generate cortical pulling on spindle poles.

To address the role of spindle shape in spindle rotation, we sought a mutant with more elongated spindles. Depletion of dynein, dynactin, LIN-5 or ASPM-1 results in long spindles (Ellefson and McNally, 2011; Connolly et al., 2014; Crowder et al., 2015), however, these proteins are implicated in dynein-dependent force generation. Katanin is a microtubule-severing protein composed of two subunits encoded by MEI-1 and MEI-2 in *C. elegans*. While, strong loss-of-function katanin mutants assemble meiotic spindles with no ASPM-1 labelled spindle poles (McNally and McNally, 2011; Connolly et al., 2014), the partial loss-of-function mutant, *mei-2(ct98)*, which has reduced microtubule-severing activity (McNally et al., 2014), assembles elongated bipolar spindles with rotation defects (McNally et al., 2006). Here we tested whether this mutant could be used to separate the effects of spindle shape from cortical force generation.

## Results

### *mei-2(ct98)* spindles frequently fail to rotate but are not defective in dynein-dependent cortical pulling

Whereas 100% (n=22) of control meiotic spindles rotated to a stable perpendicular orientation (60-90° to the tangent of the curve of the cortex) (Fig. 1A), 63% (n=30) of *mei-2(ct98)* spindles failed to rotate and anaphase proceeded on parallel or diagonal spindles (15-60° to the tangent of the curve of the cortex) (Fig. 1B, 1C). In some cases (11/30 embryos), the *mei-2(ct98)* spindle started to rotate but stalled at a shallow angle with one pole touching the cortex (Fig. 1B). In more extreme cases (8/30 embryos), anaphase proceeded on spindles that had both poles near the cortex (Fig. 1C). Two polar bodies frequently started to form over both separated masses of chromosomes on these completely parallel spindles but then regressed. The frequency of completely parallel anaphases corresponds closely with embryonic lethality in this strain (24.2% n= 347; Fisher’s exact test p = .82). Because there is some subjectivity in measuring the angle of a spindle relative to an irregular ellipsoid surface, we sought a more objective criterion for correct rotation. *C. elegans* meiotic spindles have 6 regularly spaced bivalents so that 3 – 4 bivalents are visible in a mid focal plane. In 38/40 control spindles, 3 or more homologous chromosomes merged with the cortex during anaphase (Fig. 1A, 1:45; Fig. 3F, 4:00). In contrast, in the majority (30/40) of *mei-2(ct98)* spindles, fewer than 3 homologous chromosomes merged with the cortex (Fig. 1B, 1:45).

**Figure 1.**
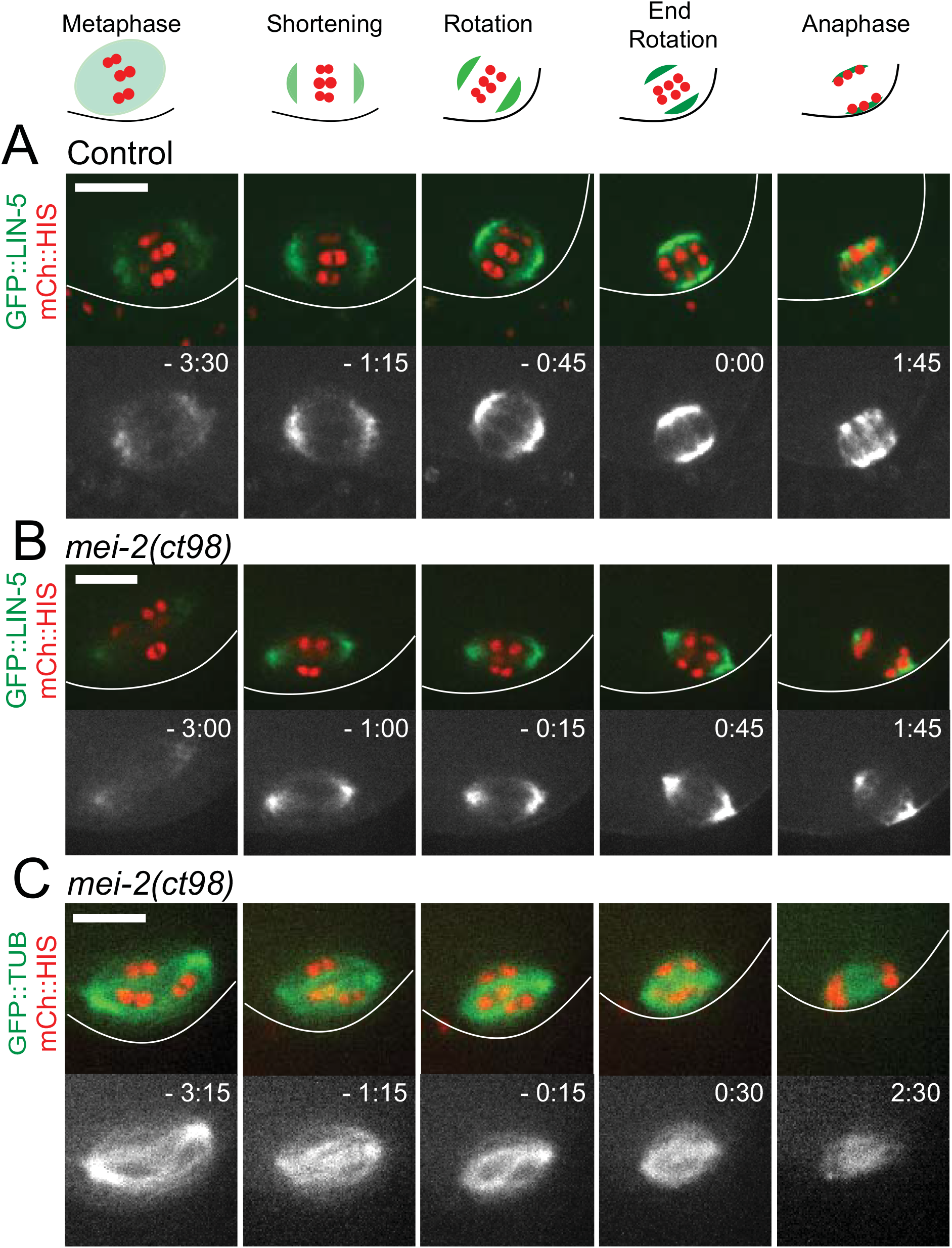
*mei-2(ct98)* meiotic spindles are defective in APC-dependent rotation. (A) Time-lapse images of a control embryo expressing GFP::LIN-5 and mCherry::Histone show the progression of the spindle from metaphase to the end of anaphase A. (B) A spindle in a *mei-2(ct98)* embryo expressing GFP::LIN-5 and mCherry::Histone undergoes a partial rotation. (C) A spindle in a *mei-2(ct98)* embryo expressing GFP::Tubulin and mCherry::Histone fails to rotate and remains parallel to the cortex during anaphase. Images were captured every 15 seconds. Time 0 is 15 seconds prior to the observed start of chromosome separation. This corresponds approximately to the start of rotation in control embryos. Bars, 6 μm.

*mei-2(ct98)* rotation failure could be due to defective dynein-dependent cortical pulling. To test this possibility, we compared the velocities of late spindle movement in 18 time-lapse sequences of *kca-1(RNAi)* embryos with those in 18 time-lapse sequences of *mei-2(ct98) kca-1(RNAi)* embryos. Whereas velocities could be determined from all of these sequences, spindle orientation could only be unambiguously scored in a subset, either because of the angle relative to the imaging plane or due to spherical aberration for deeper spindles. In embryos depleted of KCA-1, metaphase spindles are far from the cortex until spindle shortening initiates. As spindles shorten, they move toward the cortex in an APC- (Yang et al., 2005) and dynein-dependent (Ellefson and McNally, 2009) manner (Fig. 2A). In the majority of *kca-1(RNAi)* time-lapse sequences (12/13), this movement was initially a sideways “late translocation” followed by rotation to a perpendicular orientation at the cortex. *kca-1(RNAi)* spindles underwent late translocation at 2.83 +/− .30 μm/min (n=18) (Fig. 2B, D) and *mei-2(ct98); kca-1(RNAi)* spindles underwent late translocation (15/16 sideways) at 2.67 +/− .18 μ m/min (n=18) (Fig. 2C, D). Of the *mei-2(ct98) kca-1(RNAi)* spindles which translocated, 6/12 also rotated upon reaching the cortex, while 6/12 remained in a parallel orientation to the cortex during anaphase. The sideways spindle orientation and identical late translocation velocities between *kca-1(RNAi)* and *mei-2(ct98); kca-1(RNAi)* suggested that dynein dependent cortical pulling is acting on both spindle poles and that the differences in viscous drag on long vs short spindles are insignificant relative to pulling by multiple dynein motors. In addition, these results indicated that rotation failure in *mei-2(ct98)* embryos is not due to a lack of dynein-dependent cortical pulling.

**Figure 2.**
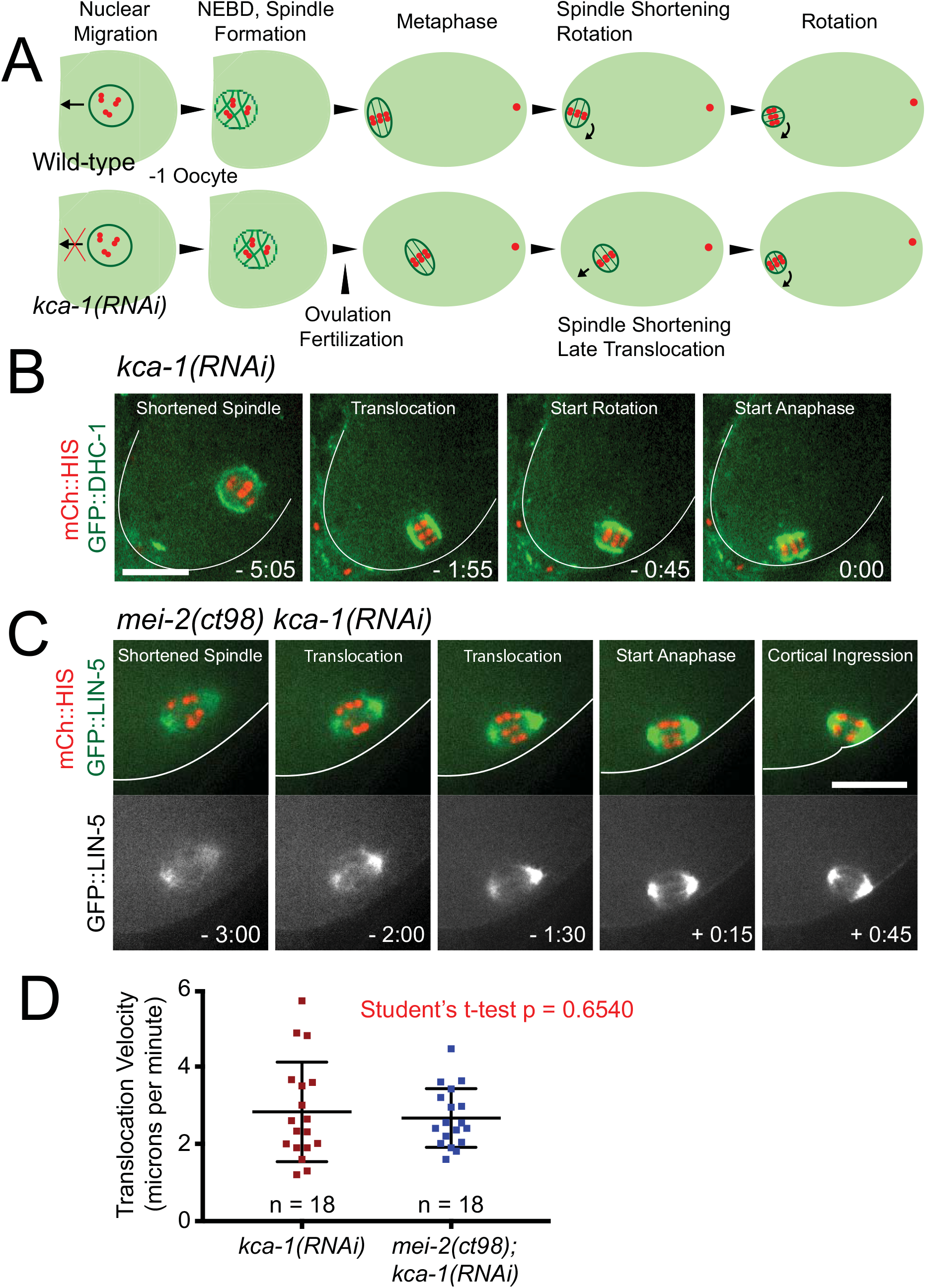
Dynein-dependent pulling forces are wild-type in *mei-2(ct98)* embryos. (A) Cartoon depiction of nuclear and spindle movements in wild-type and KCA-1-depleted worms. In wild-type oocytes, the nucleus migrates to the cortex immediately prior to nuclear envelope breakdown. The MI spindle forms and remains somewhat parallel to the cortex until APC activation and rotation. In a KCA-1-depleted embryo, nuclear migration fails and the spindle undergoes a late, dynein-dependent translocation to the cortex prior to rotation. (B) Time-lapse images of a *kca-1(RNAi)* embryo correspond to the spindle stages shown in (A). (C) In a *mei-2(ct98); kca-1(RNAi)* embryo the spindle undergoes a late translocation followed by a partial rotation. (D) Late translocation velocities are similar in *kca-1(RNAi)* embryos (2.83 +/- 0.30 s.e.m., n=18), and *mei-2(ct98); kca-1(RNAi)* embryos (2.67 +/- 0.18 s.e.m, n=18). Times in (B) and (C) are relative to the start of chromosome separation. Bars, 10μm.

As a second assay for dynein-dependent cortical pulling, we monitored endogenously tagged p50 dynamitin (GFP::DNC-2), a conserved regulator of cortical dynein required for spindle rotation (Crowder et at., 2015). In 11/11 control (Fig. 3A) and 18/18 *mei-2(ct98)* embryos (Fig. 3B), the fluorescence intensity of GFP::DNC-2 increased during APC-activated spindle shortening, at both spindle poles and the embryo cortex. In both control and *mei-2(ct98)* embryos, the fluorescence intensity of cortical GFP::DNC-2 (Fig. 3C) and cortical GFP::LIN-5 (Fig. 3D,E) increased significantly after initial contact of a pole with the cortex. Spindle pole LIN-5 appeared to merge with cortical LIN-5 as the cortex brightened (Fig. 3D, E) suggesting a possible positive feedback loop that might lock the spindle pole onto the cortex. This positive feedback loop was further suggested by control time-lapse sequences showing that chromosomes are pressed into the plasma membrane shortly after rotation (Fig. 3F). These results further indicated that *mei-2(ct98)* embryos have normal levels of dynein-dependent cortical pulling but that both poles frequently engage in cortical pulling.

**Figure 3.**
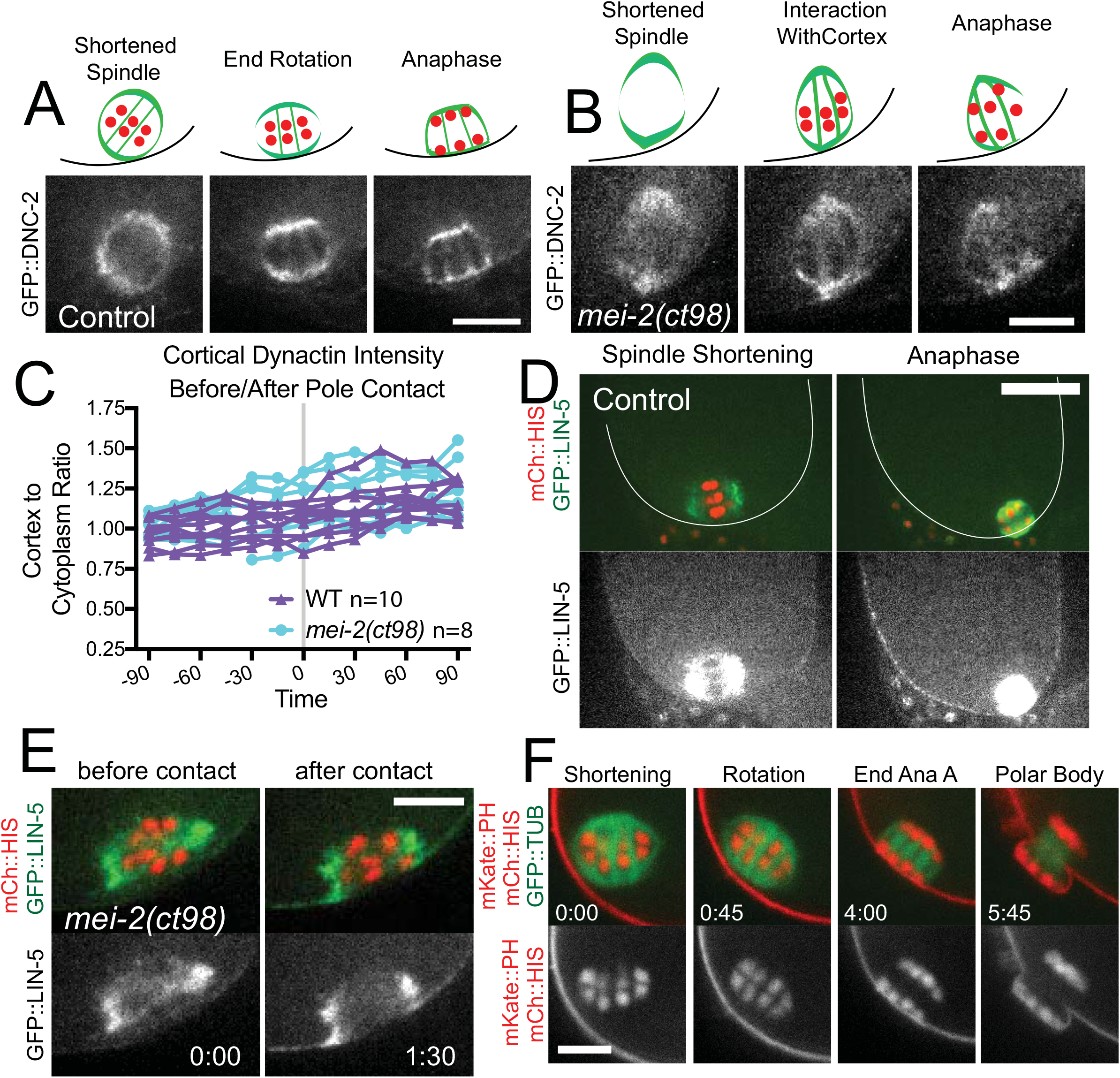
Endogenously tagged dynein regulators label spindle poles and cortices in control and *mei-2(ct98)* embryos. (A, B). GFP::DNC-2 (p50 dynamitin) increases in intensity as spindles shorten and rotate in control (A) and *mei-2(ct98)* (B) embryos. (C) Cortical GFP:DNC-2 far from the spindle increased after contact of a pole with the cortex in both control and *mei-2(ct98)* embryos. (D,E) Cortical GFP::LIN-5 increased after spindle pole contact with the cortex in control (D) and *mei-2(ct98)* (E) embryos. (F) The spindle is pressed tightly into the plasma membrane after rotation. Bars (A, B) 5 μm, (D) 10 μm, (E, F) 4 μm.

### Both spindle and pole shape correlate with rotation failure

We previously reported a correlation between wild-type spindle rotation and an axial ratio of 1.0 at the time of rotation (Crowder et al., 2015). We therefore looked for a correlation between axial ratio and rotation in *mei-2(ct98)* spindles. *mei-2(ct98)* spindles were longer than controls at metaphase and underwent shortening to variable extents. Because wild-type rotation initiates roughly 15 sec before anaphase onset, we determined the axial ratio 15 sec before anaphase onset to allow comparison of spindles that did not rotate with those that did. At this timepoint, axial ratios of *mei-2(ct98)* spindles that failed to rotate were significantly greater than those of control spindles or *mei-2(ct98)* spindles that did rotate (Fig. 4A). In addition to pre-rotation shortening, control spindles continue to shorten to a minimum axial ratio of 0.8 after rotation, before initiating anaphase B spindle elongation. The minimum axial ratios of *mei-2(ct98)* spindles that did not rotate were also greater than those of control spindles or *mei-2(ct98)* spindles that did rotate (Fig. 4B). Although the majority of spindles shortening to axial ratios of 1.5 or greater did not rotate, 3 long spindles did rotate. These spindles were already oriented at steep angles (>60°) and rotated further toward 90° whereas long *mei-2(ct98)* spindles that started at shallow angles either maintained the parallel orientation or became more parallel (Fig. 4C).

**Figure 4.**
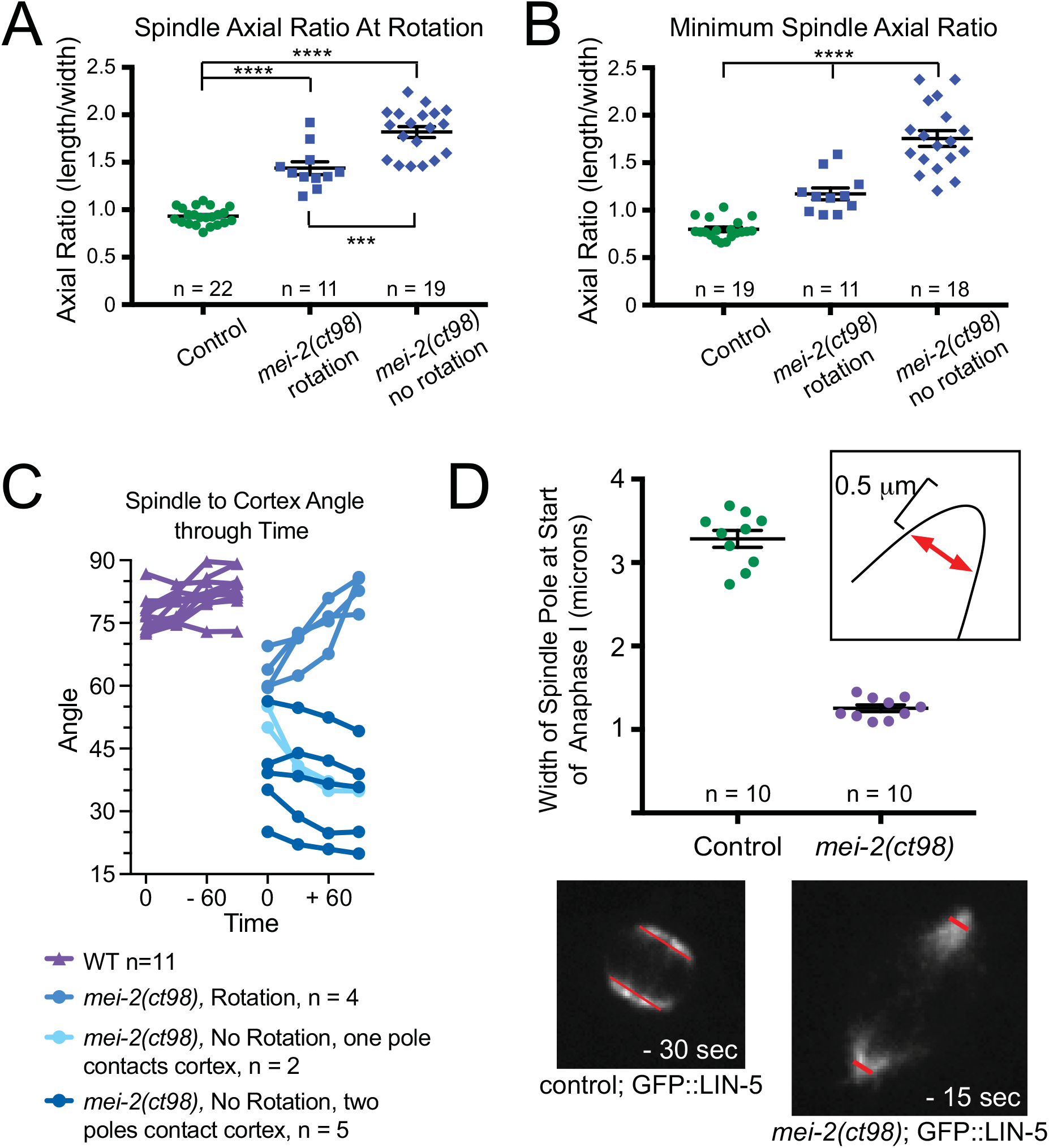
Spindle rotation is affected by both the axial ratio and the pole width. (A) Control and *mei-2(ct98)* spindles were measured 15 seconds prior to the start of chromosome segregation. Average axial ratios were: Control, 0.93 +/- 0.02 n=22; *mei-2(ct98)* which rotate, 1.44 +/- 0.07 n=11; *mei-2(ct98)* which do not rotate, 1.82 +/- 0.06 n=19. (B) Control and *mei-2(ct98)* spindles were measured at the end of anaphase A. Those *mei-2(ct98)* spindles that rotated shortened during anaphase A while the non-rotated *mei-2(ct98)* spindles did not. Control, 0.80 +/- 0.02 n=19; *mei-2(ct98)* which rotate, 1.17 +/- 0.06 n=11; *mei-2(ct98)* which do not rotate, 1.75 +/- 0.08 n=18. ***, P < 0.001; ****, P < 0.0001. (C) Spindle to cortex angle was measured in time-lapse images, captured at 15 second intervals, of embryos expressing GFP::DNC-2. t=0 is when a pole touches the cortex. (D) Spindle pole width (red arrow) was measured 0.5mm from the outermost edge of the spindle. Time in the examples is from the start of chromosome separation. Average widths are: control, 3.29 +/- 0.10 n=10; *mei-2(ct98)*, 1.25 +/- 0.04 n=10.

In addition to large axial ratios, *mei-2(ct98)* spindles had poles that remained pointed in shape throughout metaphase and anaphase (Fig. 1B; Fig. 4D), whereas control spindle poles became round as rotation started and became partially flattened during rotation so that the curve of the pole closely matched the curve of the cortex (Fig. 1B; 4D). If the width of the LIN-5 measured 0.5 μm from the tip of the spindle is used to calculate the potential surface area of contact between LIN-5 on the pole and LIN-5 on the cortex after rotation to a perpendicular orientation, control spindles would have an average of 8.5 μm^2^ of contact whereas *mei-2(ct98)* spindles would only have 1.3 μm^2^ of contact. The pointed *mei-2(ct98)* spindle poles would clearly maximize the surface area of contact between pole LIN-5 and cortical LIN-5 when one spindle pole is apposed to the cortex at a shallow angle as can be seen in Fig. 1B. Thus large axial ratios and pointed spindle poles both correlated with rotation failure of spindles that have normal dynein-dependent cortical pulling force (Fig. 2).

### A PIP2-binding plekstrin homology domain suppresses *mei-2(ct98)* spindle rotation defects by promoting spindle shortening and spindle pole flattening

In order to better image polar body formation induced by parallel anaphase spindles, we constructed a *mei-2(ct98)* strain expressing an mKate2 fusion to the PIP2-binding plekstrin homology domain of rat phospholipase C delta. Surprisingly, embryonic lethality at 25°C was significantly reduced (p=.0001) from 24% (n=347) to 14% (n=700) and the majority of embryos produced by these worms underwent perpendicular anaphase (12/14 rotated to 60° or greater; Fig. 5C). At metaphase, these PH-suppressed *mei-2(ct98)* parallel spindles had high axial ratios and pointed poles similar to those of the non PH-suppressed strain (Fig. 5C, D). However, when one pole contacted the PH at the cortex, the spindles shortened and the cortex-proximal pole flattened to allow rotation (Fig. 5C, E). Whereas the mechanism of this PH domain effect on spindle shape is not known, this result strengthens the correlation between spherical spindle shape and successful rotation to 90°.

**Figure 5.**
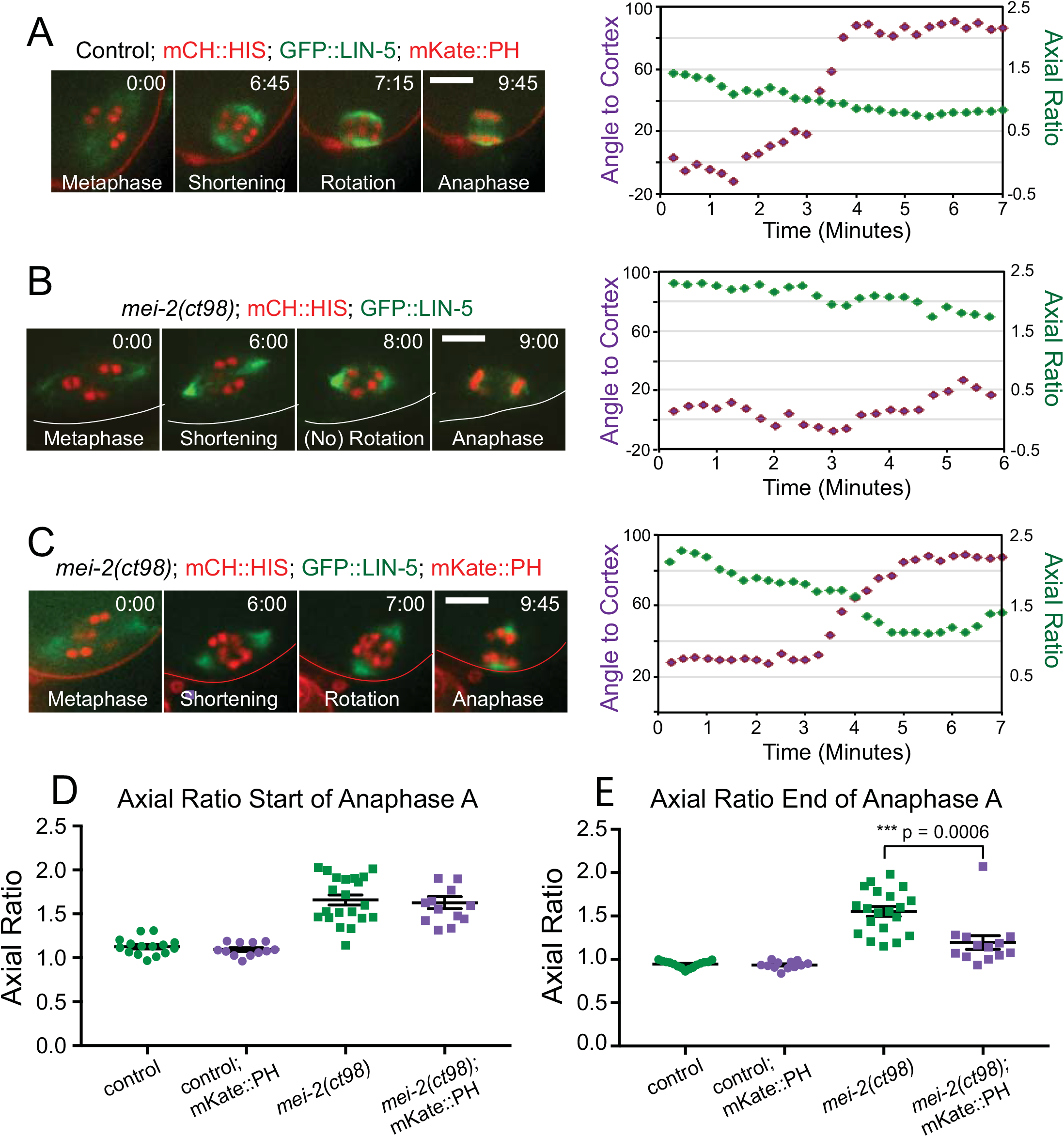
The rotation defect in *mei-2(ct98)* embryos is suppressed by an mKate::PH transgene. (A) Representative time-lapse images are shown for a control embryo expressing mCh::HIS, GFP::LIN-5 and mKate::PH. A plot of both axial ratio and spindle to cortex angle over time indicates that rotation began when the axial ratio was 1.0 and was completed within 60 seconds. (B) Representative time-lapse images from a *mei-2(ct98)* embryo expressing mCH::HIS and GFP::LIN-5 only show that no rotation occurs. All Bars, 5 μm. (C) Time-lapse images captured from a *mei-2(ct98)* embryo expressing mKate::PH as well as mCh::HIS and GFP::LIN-5 show that spindle rotation began when the axial ratio was 1.72 and continued for 90 seconds. (D) Axial ratios are shown for control and *mei-2(ct98)* spindles at the start of chromosome separation. Control, no PH: 1.13 +/- 0.03, n=15; control with PH: 1.09 +/- 0.02,n=12; *mei-2(ct98)*, no PH: 1.66 +/- 0.06, n=21; *mei-2(ct98)* with PH: 1.63 +/- 0.07, n=13. (E) Axial ratios of control and *mei-2(ct98)* spindles at the end of anaphase A. Control, no PH: 0.95 +/- 0.01, n=15; control with PH: 0.93 +/- 0.01, n=12; *mei-2(ct98)*, no PH: 1.55 +/- 0.06, n=20; *mei-2(ct98)* with PH: 1.20 +/- 0.08, n=13.

### Modeling reveals that spherical spindle shape reduces the probability of isometric cortical pulling on both spindle poles

To separate the possible role of spindle shape from other factors that change in any mutant scenario, we generated Cytosim (Nedelec and Foethke, 2007) models of rotation. In these two dimensional simulations, spindles of 4 different shapes were built as solid objects with astral microtubules extending from nucleators at the spindle poles. “Short flat” spindles had aspect ratios of 0.9 and slightly flattened poles to mimic the shape of wild-type spindles mid-rotation. “Long pointed” spindles had aspect ratios of 1.5 and pointed poles to mimic the shape of *mei-2(ct98)* spindles that fail to rotate. To distinguish between the relative importance of axial ratio vs flattened poles, imaginary “long flat” and “short pointed” spindles were also built. Astral microtubules emanated from nucleators arranged to mimic the localization of LIN-5/dynein in wild-type or *mei-2(ct98)* spindle poles and astral microtubules underwent dynamic instability. Starting with in vivo dynamic instability parameters taken from Kline-Smith and Walczak (2002), a limited parameter sweep was conducted to find conditions that allowed short flat spindles to rotate within 300 seconds, the normal time between initiation of spindle shortening, indicative of APC activation, and completion of anaphase A. Short 200 nm cortical microtubules emanated from nucleators at the cortex (green dots in Fig. 6A). Dynein complexes (red dots) in the cytoplasm bound to astral or cortical microtubules, motored toward minus ends and dissociated with a rate constant that increased with applied force. When a dynein complex contacted an astral microtubule and a cortical microtubule, the spindle was pulled toward the cortex. A stabilizing “cortical glue” (purple dots) was included so that spindles became more stably attached once close contact between the pole and the cortex was achieved. This cortical glue was intended to mimic the positive feedback loop suggested in Fig. 3. Spindles were initially perfectly parallel to and 2 μm away from the cortex at the start of each simulation, and simulations were run for 300 seconds. In nearly all cases, spindles initially moved toward the cortex in a sideways orientation and astral microtubules from both poles made pulling contacts with cortical microtubules for durations ranging from 3 – 248 seconds (Movies 1-5). The initial sideways movement of simulated spindles matched the initial sideways motion observed in 4/6 control embryos with a clearly labeled plasma membrane (Fig. 6C; see also Fig. 1A in Crowder et al., 2015). In these simulations, 100% (n=99) of short flat spindles rotated to 90° (Movie 1) whereas only 3% (n=100) of long pointed spindles rotated to angles of 60-90°, balancing on the tip of the pointed pole (Fig. 6A). 97% of long pointed spindles partially rotated but became stuck with one flat side of the pole stably apposed to the cortex (Movie 2), similar to the majority of *mei-2(ct98)* spindles (Fig. 1B). To separate axial ratio from pole shape, we ran simulations of imaginary short pointed spindles and long flat spindles (Fig. 6A; Movie 1-2). 78% (n=100) of short pointed spindles rotated to 60-90°, balancing on the point of the spindle while 22% became stuck on one flat side of a pole. Thus an axial ratio of 0.9 significantly (p<.0001 Fisher’s exact test) improved the success rate of rotation with pointed poles relative to an axial ratio of 1.5. Long flat spindles successfully rotated to 60-90° in 94% (n=100) of simulations and were stuck parallel to the cortex with isometric cortical pulling on both poles in only 6% of simulations. Thus flat poles significantly (p<.0001 Fisher’s exact test) improved the efficiency of rotation to 90° relative to pointed spindles even with a constant axial ratio of 1.5.

**Figure 6.**
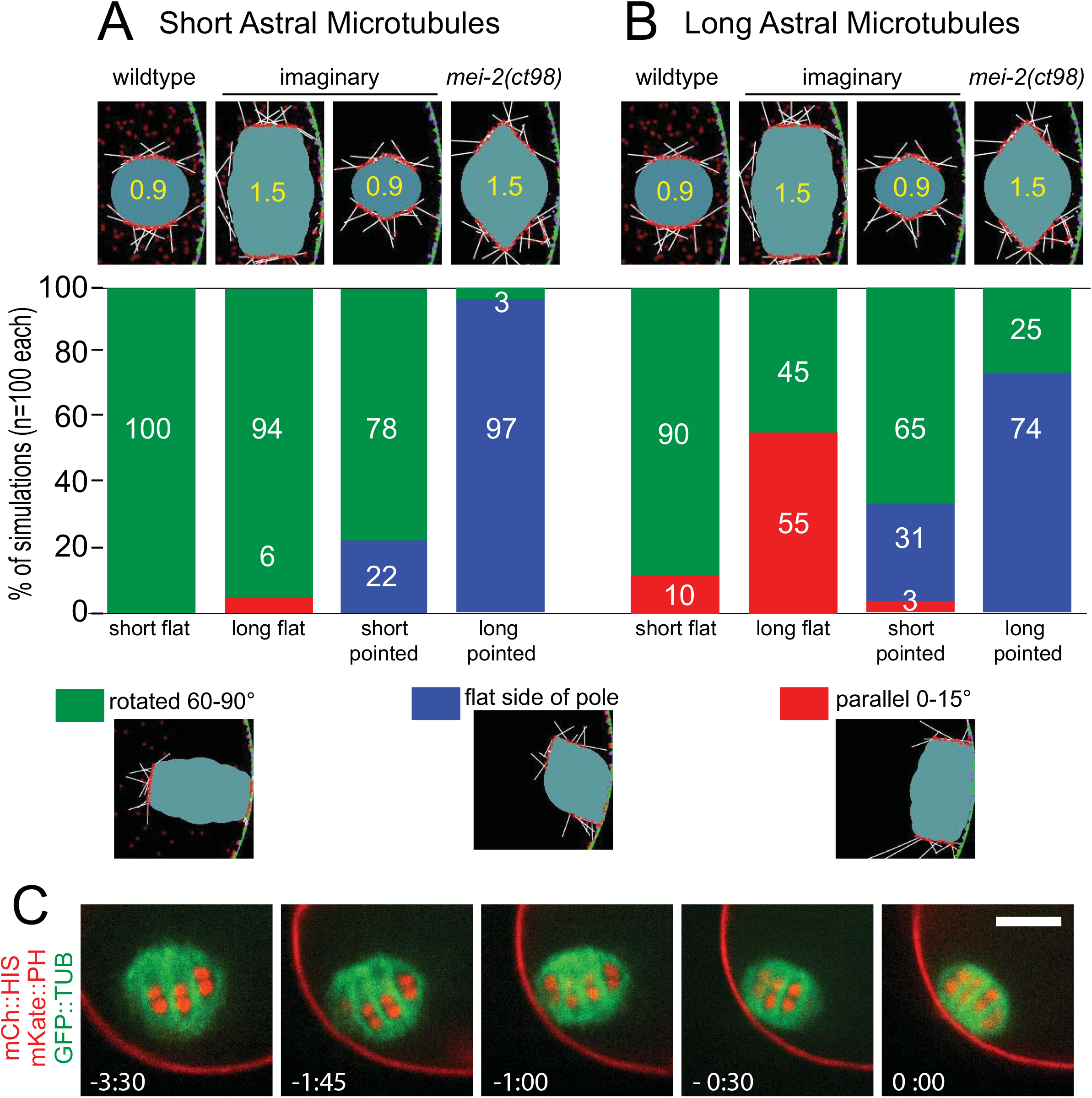
Simulations reveal that both short axial ratios and flattened spindle poles contribute to successful rotation to 90°. A) and B) Frequency of successful rotation to 90° during 300 second Cytosim simulations with spindles of different shapes. n=100 simulations for each of 8 conditions. C) Time-lapse sequence of a real rotation event showing that the spindle initially moves sideways toward the cortex before rotating. Bar = 5 μm.

We next considered why long pointed spindles never became stuck in a parallel orientation in our simulations. From visual inspection of these partially rotated spindles (Fig. 6A bottom; Movie 2), it is obvious that astral microtubules are not long enough to reach from the cortex distal pole to the cortex and that to maximize pole-cortex contact on both poles, the spindle would have to bend. To address the role of astral microtubule length, we ran the same simulations but with the polymerization rate tripled from .06 μm/s to 0.18 μm/s (Fig. 6B; Movie 3-4). This polymerization rate is faster than that reported in cultured mammalian cells but still slower than that reported in *C. elegans* mitotic embryos (Srayko et al., 2005). The resulting longer astral microtubules did not cause long pointed spindles to remain parallel. In contrast, longer astral microtubules completely blocked rotation in 10% of short flat spindles and 55% of long flat spindles by generating isometric cortical pulling on both poles. The rotation-promoting effect of a 0.9 axial ratio vs a 1.5 axial ratio for flat-poled spindles with longer astral microtubules was extremely significant (p<.0001). The simulations with longer astral microtubules solidified the conclusion that both short axial ratio and flat poles promote rotation, however, astral microtubule length does not appear to explain why simulations of long pointed spindles do not get stuck in a parallel orientation.

To compare the time needed to reach a 60-90° perpendicular spindle orientation under different simulated conditions using ANOVA, we set the rotation time for those spindles that did not actually achieve a perpendicular orientation to 310 seconds (Fig. 7). Long spindles took significantly longer to rotate than short spindles with the same pole shape and pointed spindles took significantly longer to rotate than flat-poled spindles with the same axial ratio. This analysis again indicated that both short axial ratios and flattened poles can promote spindle rotation, but again did not explain why simulated long pointed spindles did not get stuck in a parallel orientation.

**Figure 7.**
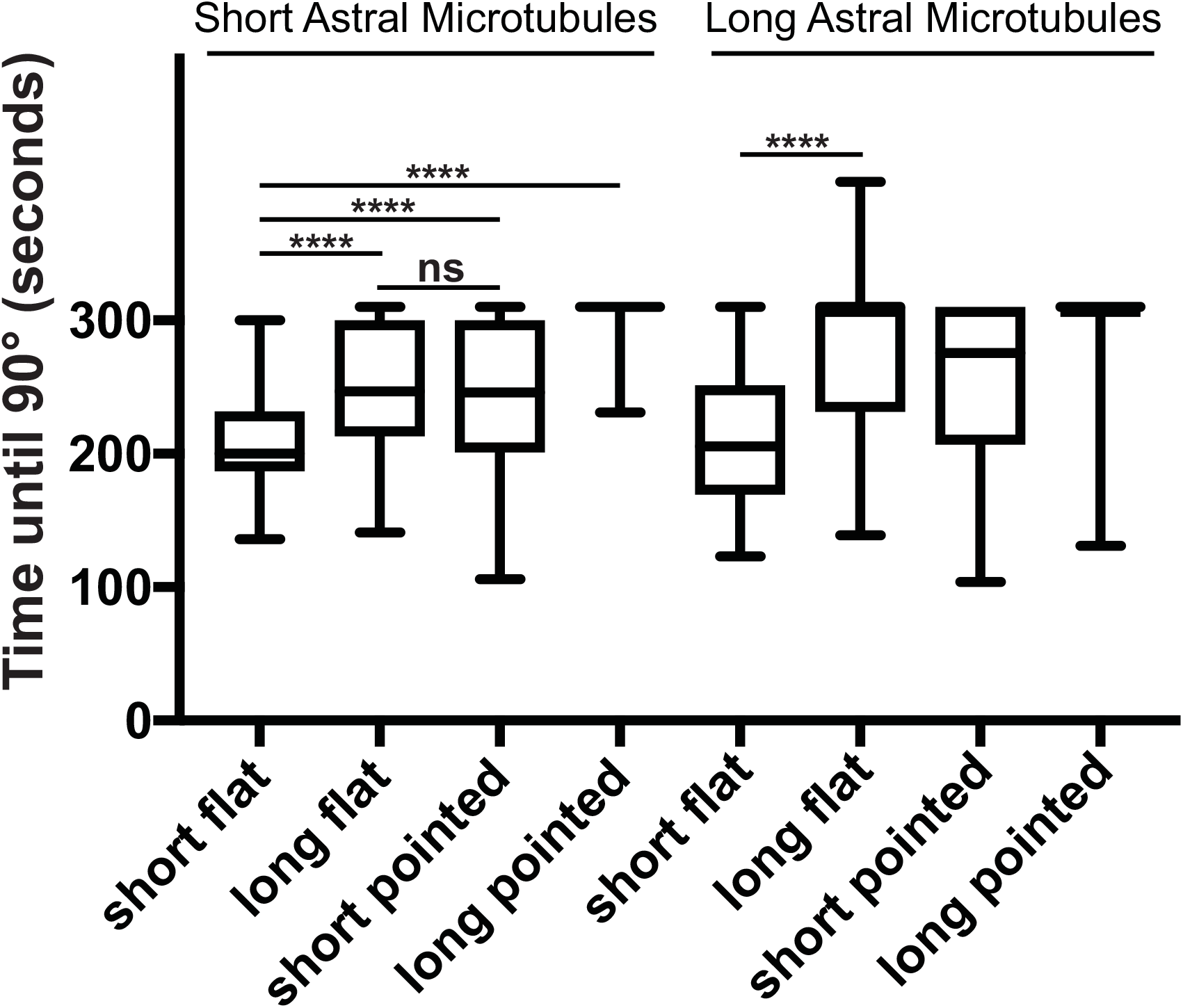
Both short axial ratio and flat spindle poles decrease the time required to rotate to 90° in Cytosim simulations. Spindles that did not rotate to 90° during the 300 sec simulation were assigned a time of 310 sec. n=100 simulations for each of 8 conditions. **** indicates p<.0001 by ANOVA.

### *mei-2(ct98)* spindles have a low density of microtubules and bend to allow cortical capture of both spindle poles

Time-lapse imaging (Fig. 1A and B), as well as simulation results that include a “cortical glue” to mimic the positive feedback loop suggested by Fig. 3, both indicated that the surface area of contact between pole LIN-5/dynein and cortical LIN-5/dynein is maximized during rotation. Simulated long pointed spindles cannot maximize contact between both poles and the cortex because of their fixed shape. As shown in the time-lapse sequence in Fig. 8A, *mei-2(ct98)* spindles bent back and forth during both metaphase and anaphase B. *mei-2(ct98)* spindles had a reduced density of both EBP-2::mKate, which tracks growing plus ends of microtubules (Fig. 8B, C), and GFP::tubulin (Fig. 8D; see also McNally et al., 2014). The combination of large axial ratio with reduced density of microtubules likely allows spindle bending, which in turn allows both poles to come into close enough proximity for stabilized cortical capture.

**Figure 8.**
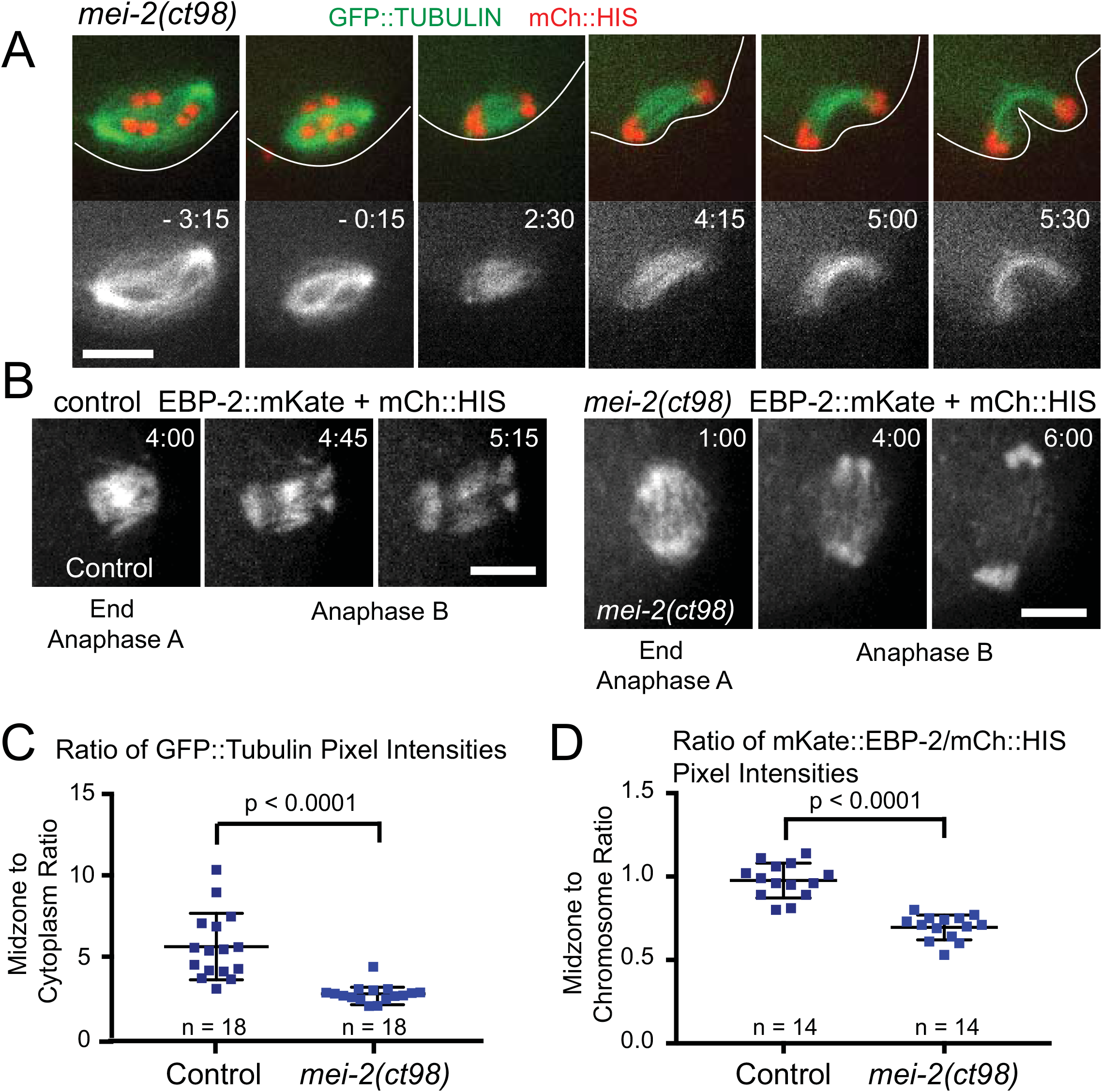
Both spindle poles in *mei-2(ct98)* spindles can interact with the cortex because low MT and EBP-2 density in the midzone allows them to bend. (A) Time-lapse images of a *mei-2(ct98)* embryo expressing GFP::tubulin and mCh::HIS shows a spindle which is parallel to the cortex during anaphase and which bends as the cortex invaginates. Time is relative to the start of chromosome separation. Bar, 6μm. (B) Time-lapse images of control and *mei-2(ct98)* embryos expressing EBP-2::mKate2 and mCherry::histone. Bars, 4μm. (C) Mean GFP::tubulin pixel intensity for the midzone and the cytoplasm was measured when the chromosomes were 4-5μm apart and background subtracted. The average ratio for the control was 5.84 +/- 0.63 s.e.m.; the average *mei-2(ct98)* ratio was 2.84 +/- 0.21 s.e.m. (D) Mean EBP-2::mKate2 pixel intensity in the midzone and mean mCh::HIS pixel intensity were measured when the chromosomes were 4-5μm apart. The average ratio for the control was 0.98 +/- 0.03 s.e.m.; the average ratio for *mei-2(ct98)* was 0.69 +/- 0.02 s.e.m.

### F-actin and ER might resist dynein-mediated cortical pulling

Simulations of short vs long flat-poled spindles with long astral microtubules (Movie 3) indicated that spindles with a long axial ratio can remain parallel with both poles engaged in cortical pulling for a longer period of time than spindles with a short axial ratio (Fig. 7). The reason for this difference, however, was not clear. The difference might be caused by shape-dependent, steric masking of astral microtubules during rotation or it might be caused by shape-dependent differences in viscous drag. In most simulations, astral microtubules from both poles initially engaged the cortex, then one pole lost cortical attachments before pivoting away from the cortex. If long spindles pivot more slowly due to higher translational and rotational drag, this might allow more time for astral microtubules from the departing pole to recapture the cortex. Our Cytosim models used a cytoplasmic viscosity of 1 Pascal sec, determined from microrheology of 100 nm PEG-coated beads in *C. elegans* embryos (Daniels et al., 2006). Both translational drag that depends on the radius of the spheres used to build the spindles and rotational drag that depends on the distance between spheres were incorporated into our Cytosim models. To determine whether viscosity might have an effect on spindle rotation, we ran simulations of short flat vs long flat spindles with long astral microtubules with a 10-fold reduction in viscosity (Movie 5). Rotation was completely rescued (20/20) for long spindles and this increased frequency of successful rotation by reducing viscosity (45% n=100 to 100% n=20) was significant (p<.0001 Fisher’s Exact test) and the time until reaching 90° was not significantly different between short and long spindles at low viscosity (short: 188 +/- 15 sec, n=11; long 164 +/- 12 sec, n=20; p=.24 t test). In contrast, rotation of short flat spindles was relatively unaffected by reducing viscosity (20% failure n=100 vs 20% failure n=20; time to rotation high viscosity: 214 +/- 6 sec n=99 vs low viscosity: 188+/-15 sec n=20, p=.08 t test).

F-actin can resist movement in a size and shape-dependent manner that is analogous but distinct from viscosity (Feric and Brangwynne, 2013). To experimentally test whether F-actin generates any measurable opposition to cortical pulling, we depleted profilin by RNAi to disassemble F-actin, and then analyzed its effects on spindle translocation in the kinesin-1 heavy chain mutant, *unc-116(f130)*. This Ala236Thr point mutation causes a temperature sensitive maternal effect lethal neomorphic phenotype with the meiotic spindle completing meiosis in the middle of the embryo. The metaphase spindle is further from the cortex than in *unc-116(RNAi)* or *kca-1(RNAi)* but can move to the cortex if dynein is hyper-activated by CDK1 inhibition (Ellefson and McNally, 2011). Movement of the spindle to the cortex was also induced by depolymerizing F-actin by profilin (RNAi) (Fig. 9A-B). This result indicated that F-actin, which forms a deep cytoplasmic meshwork in meiotic embryos (Panzica et al., 2017), can oppose dynein-dependent cortical pulling. To investigate whether the endoplasmic reticulum (ER) might cause drag on meiotic spindle rotation, we conducted time-lapse imaging of a strain expressing GFP::signal peptidase. In 8/8 time-lapse sequences of control worms, the ER formed a reticulum that interdigitated with the metaphase spindle and formed dense accumulations at spindle poles during metaphase and the early stages of rotation (Fig. 10A; Poteryaev et al., 2005). The morphology of the ER changed to become more vesicular during spindle rotation but the ER remained around the spindle and between the spindle and cortex during rotation. A similar transition in the ER occurred at the onset of late translocation in *kca-1(RNAi)* embryos, however, ER was associated with the spindle throughout late translocation (Fig. 10B). Reticular ER did not move with kinesin-driven cytoplasmic streaming (Fig. 10C) whereas the vesicular ER moved at an average velocity of 0.23 μm/sec (range 0.1 – 0.6 μm/s; n=5 embryos) similar to the velocity of kinesin-driven yolk granule movement (McNally et al; 2010) as previously reported (Kimura et al., 2017). Because reticular ER has the mechanical strength to resist cytoplasmic streaming, it could generate resistance to dynein-driven cortical pulling.

**Figure 9.**
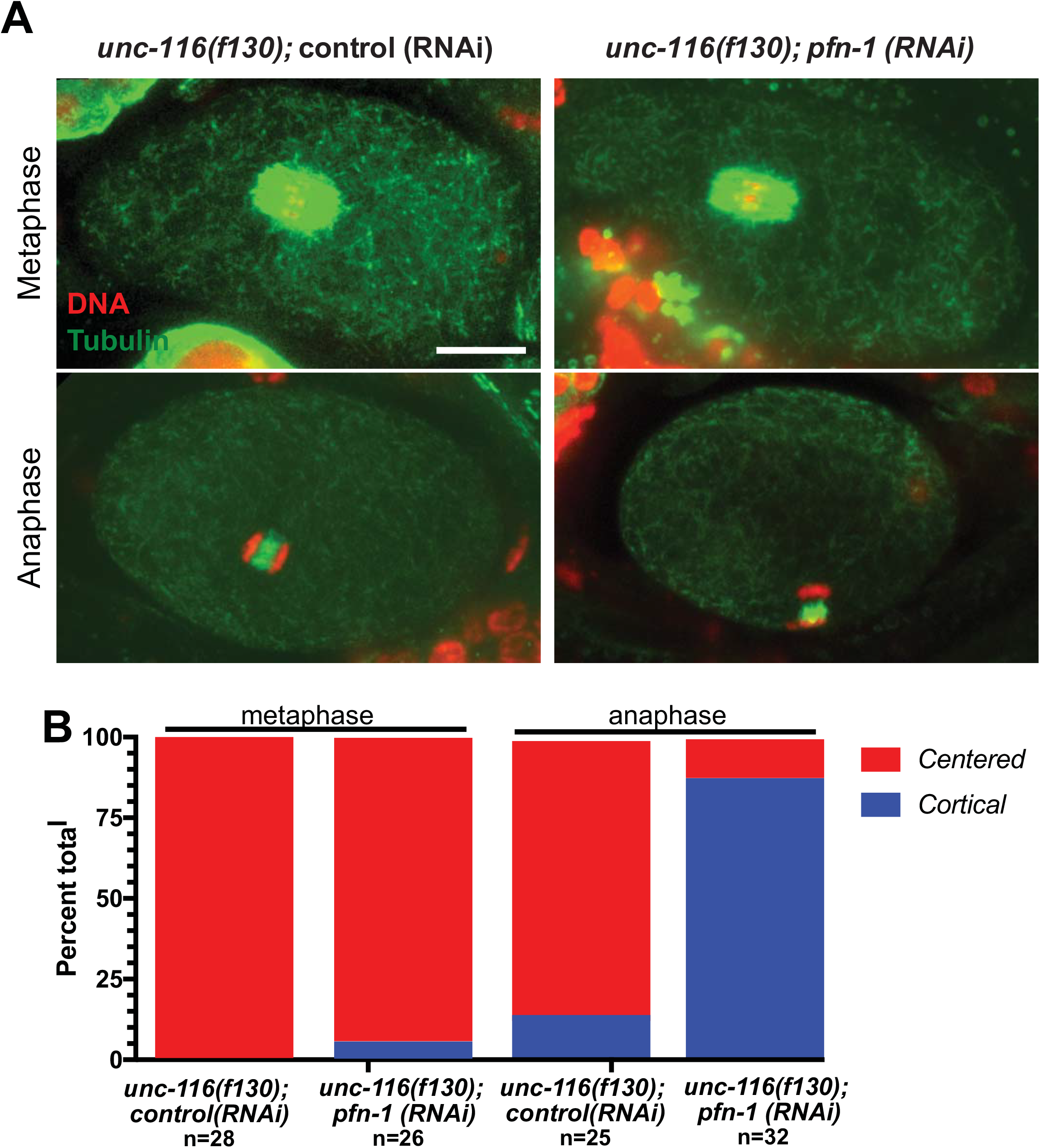
Depletion of F-actin rescues late spindle translocation in *unc-116(f130)* meiotic embryos. (A) Representative images of embryos grown at 25°C, fixed and stained with anti-tubulin antibodies. Bar, 10 μm. (B) Quantification of the frequency of centered or cortical spindle position at metaphase and anaphase.

**Figure 10.**
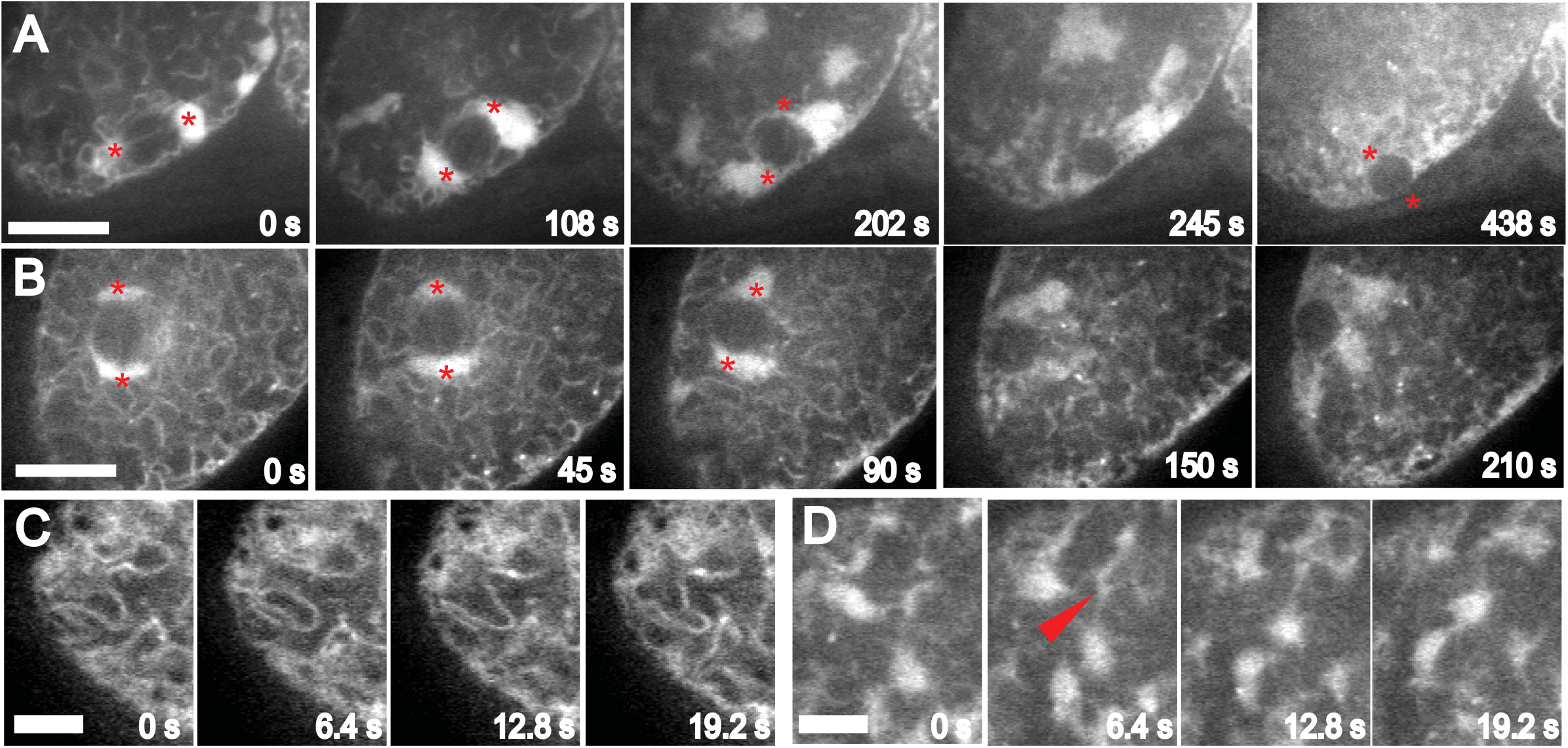
The ER as a potential restraint on cortical pulling. A-D. Time-lapse images of the ER labelled with GFP:signal peptidase. A. Control and B. *kca-1(RNAi)* show sheet-like ER during metaphase changing to vesicular ER during spindle shortening and rotation or late translocation. Asterisks indicate the position of spindle poles. C-D. Higher frame rate images showing that sheet-like ER (C) is stationary while vesicular ER (D) is moving. Arrowhead indicates the direction of movement. Bar in A, B = 10 μm. Bar in C,D = 5 μm.

## Discussion

Our combined experimental and simulation results strongly support a model in which both short axial ratio (1-0.9) and slightly flattened spindle poles promote meiotic spindle rotation to a perpendicular orientation relative to the cortex. This perpendicular spindle orientation during anaphase is extremely conserved across animal phyla. The fact that the frequency of parallel anaphase matches the frequency of unhatched embryos for *mei-2(ct98)* supports the idea that perpendicular spindle orientation facilitates accurate extrusion of chromosomes into polar bodies. Short axial ratio with slightly flattened poles is conserved in other species with acentriolar female meiotic spindles such as humans (Holubcova et al., 2015) and even in species with centriole-bearing female meiotic spindles such as *Spisula solidissima* (McNally et al., 2016). Drosophila female meiotic spindles are naturally long and pointed during early metaphase, then shorten and round before rotating (Endow and Komma, 1997). Work on mouse oocytes has focused on the idea that meiotic spindles are positioned by an actin-driven and microtubule-independent mechanism (Mogessie et al., 2018), however, low doses of microtubule depolymerizing drugs inhibit meiosis II spindle rotation in mice (Ai et al., 2008a) and rats (Ai et al., 2008b). Thus it is possible that cortical pulling on astral microtubules contributes to actin-driven mouse spindle rotation and it is also possible that spindle shape promotes cortical pulling by acto-myosin on one spindle pole (Schuh and Ellenberg, 2008).

Our combined experimental and modeling results indicated that short spindles with slightly flattened poles promote accurate spindle rotation through three mechanisms. First, the round but slightly flattened shape of wild-type spindle poles during rotation maximizes the surface area of contact between pole dynein and cortical dynein. The positive feedback loop suggested by the increase in cortical dynactin after pole contact would stabilize wild-type spindles in a perpendicular orientation and *mei-2(ct98)* spindles in a diagonal orientation. Second, short spindles are likely more rigid than long spindles and thus do not bend like *mei-2(ct98)* spindles. Spindle bending makes it possible for both spindle poles to make close stable contact with the cortex. Third, the reduced viscous drag on a short spindle allows one pole to escape the cortex quickly so that astral microtubules from the escaping pole do not recapture the cortex. These three mechanisms are equally relevant for capture of spindle pole dynein by microtubule plus ends emanating from the cortex as they are for capture of cortical dynein by astral microtubules.

The finding that long *mei-2(ct98) kca-1(RNAi)* spindles undergo sideways late translocation at the same velocity as short *kca-1(RNAi)* spindles might seem contradictory with the finding that actin depolymerization facilitates late translocation of *unc-116(f130)* spindles. The first result suggests that shape-dependent drag is insignificant whereas the second results suggests that F-actin might confer enough resistance to cortical pulling to block translocation across longer distances. We previously calculated that shape-dependent differences in viscous drag would be small relative to the force of a single dynein molecule (Crowder et al., 2015). The observed parallel spindle orientation during late translocation in *kca-1(RNAi)* embryos, as well as during early stages of rotation in control embryos, suggest that this movement is mediated by multiple dynein motors pulling on both poles. During this early stage of pulling on both poles, picoNewtons of drag would be much less than the picoNewtons of pulling. In our simulations, rotation initiates when one pole loses cortical pulling attachments. At this point the competition would not be between picoNewtons of dynein pulling and picoNewtons of drag. Instead, the competition would be between the velocity of spindle pivoting and the probability of an astral microtubule recapturing the cortex through dynamic instability.

Parameters other than drag could also contribute to our results. If cortical microtubules are longer in *mei-2(ct98) kca-1(RNAi)* embryos, as reported for *mei-1(RNAi)* embryos (Kimura et al., 2017), then late translocation might be mediated by more microtubules extending between spindle poles and the cortex than in *kca-1(RNAi)* embryos. Likewise, recent work has shown that actin meshworks can inhibit microtubule polymerization in *Xenopus* extracts (Colin et al., 2018). Therefore the promotion of late translocation by actin depolymerization in *unc-116(f130)* embryos might be through an increase in the length of astral or cortical microtubules.

Understanding of the molecular mechanism of rotation would be improved by direct observation of the force-generating microtubule connections between the spindle and cortex. The localization of ASPM-1, LIN-5, dynactin and dynein to both the spindle pole and the cortex, as well as the presence of numerous cortical microtubules thought to drive cytoplasmic streaming and yolk granule packing (McNally et al., 2010), make it difficult to identify the relevant force-generating microtubules. Pursuing a 3D model of rotation in which spindles are continuously shortening and clarification of the contribution of viscous drag from both experiment and modeling will also be essential.

## Materials and Methods

### C. elegans strains

The following strains were used in this study: FM125: ruIs57[pie-1p::GFP::tubulin + unc-119(+)]V; itIs37[pie-1p::mCherry:H2B::pie-1 3’UTR + unc-119(+)] IV. FM13: mei-2(ct98) I; ruIs57[pie-1p::GFP::tubulin + unc-119(+)]V; itIs37[pie-1p::mCherry:H2B::pie-1 3’UTR + unc-119(+)] IV. FM562: lin-5(he244[egfp::lin-5] II; itIs37[pie-1p::mCherry:H2B::pie-1 3’UTR + unc-119(+)] IV. FM582: mei-2(ct98) I; lin-5(he244[egfp::lin-5] II; itIs37[pie-1p::mCherry:H2B::pie-1 3’UTR + unc-119(+)] IV. FM485: lin-5(he244[egfp::lin-5] II; is[mex-5p::mCherry::H2B + mKate2::PH inserted in K03H6.5]IV. FM583: mei-2(ct98) I; lin-5(he244[egfp::lin-5] II; Is[mex-5p::mCherry::H2B + mKate2::PH inserted in K03H6.5]IV. FM461: cpIs54[mex-5p::mKate::PLC(delta)PH(A735T)::tbb-2 3’UTR + unc-119(+)] II; itIs37[pie-1p::mCherry:H2B::pie-1 3’UTR + unc-119(+)] IV; ruIs57[pie-1p::GFP::tubulin + unc-119(+)]V. EU1561: orIs17 [dhc-1::GFP::DHC-1, unc-119(+)]; itIs37 [unc-119(+) pie-1::mCherry::H2B] IV. FM460: prtSi122[pRG629; mex-5p::ebp-2::mKate2::tbb-2 3’UTR + unc-119(+)] II; dnc-2[prt42(N-terminal 3XFLAG::GFP)] III; itIs37[pie-1p::mCherry:H2B::pie-1 3’UTR + unc-119(+)] IV. FM462: mei-2(ct98) I; prtSi122[pRG629; mex-5p::ebp-2::mKate2::tbb-2 3’UTR + unc-119(+)] II; dnc-2[prt42(N-terminal 3XFLAG::GFP)] III; itIs37[pie-1p::mCherry:H2B::pie-1 3’UTR + unc-119(+)] IV. HR399: unc116(f130) unc-36(e251) III WH327: ojIs23(GFP::SP12).

### *In utero* live imaging

L4 larvae were incubated overnight at 25°C, then adult hermaphrodites were anesthetized with tricaine/tetramisole as described (Kirby et al., 1990; McCarter et al., 1999) and gently mounted between a coverslip and a thin 3% agarose pad on a slide. Time lapse images were acquired at 15 sec intervals with a Solamere spinning disk confocal equipped with a Yokogawa CSU10, Hammamatsu ORCA FLASH 4.0 CMOS detector, Olympus 100X/1.35 objective and MicroManager software control.

### Fixed immunofluorescence

*C. elegans* meiotic embryos were extruded from hermaphrodites in 0.8× egg buffer by gently squishing of worms between coverslip and slide, flash frozen in liquid N2, permeabilized by removing the coverslip, and then fixed in cold methanol before staining with antibodies and DAPI. Primary antibodies used in this work were mouse monoclonal anti-tubulin (DM1α; Sigma-Aldrich). Secondary antibodies used were Alexa Fluor 594 anti–mouse (Molecular Probes). Complete z stacks were captured for each meiotic embryo with an inverted microscope (Olympus IX81) equipped with a 60× PlanApo 1.42 objective, a disk-scanning unit (Olympus), and a Hammamatsu ORCA Flash 4.0 CMOS detector and controlled with MicroManager software..

### RNAi

RNAi depletion of KCA-1 and PFN-1 was carried out by feeding HT115 bacteria transformed with L4440-based plasmids (Kamath et al., 2003).

### Spindle rotation simulations

The complete models are included in Online Supplemental Material. Our agent-based simulations of spindle rotation were run on Cytosim, a stochastic physics simulator specializing in cytoskeletal cellular processes. Simulations are set within a 2-dimensional representation of a *C. elegans* zygote built as a capsule (rectangle with hemispheres capping off two opposing sides) with hemisphere radii of 15 microns and rectangular length of 10 microns. Viscosity within the cell is set to 1 Pa.s, approximately the viscosity of an embryo, and temperature is set to 25C. Within this environment steric interactions are enabled, ensuring that solid objects including the spindle and microtubules cannot occupy the same space. Two different microtubule types are simulated. First, cortical microtubules nucleating and remaining stable in length at 200 nanometers. Astral microtubules are nucleated from spindle poles and exhibit dynamic instability, reaching lengths of up to a few microns before catastrophe and rescue. At the onset of simulation the spindles are built as solid objects, made up of several solid bead-like structures to approximate the shape of a spindle that is either a) shortened with flat poles, b) shortened with pointed poles, c) long with flat poles, or d) long with pointed poles. Spindles are assembled so that their pole-to-pole axis is tangential to the hemisphere center, these are placed in this orientation approximately 1.5-2 microns away from the edge of the cell, perfectly centered to the hemisphere. Spindle poles are defined in the simulation by a series of points that act as microtubule nucleators, the spatial positions of these points are user-defined in relation to the origin of the spindle solid object, ensuring that as the spindle moves in simulation space the spindle poles move with it. The cell-space edge proximal to the spindle is seeded with 140 cortical microtubule nucleators which are fixed in space in the shape of the cell boundary. This same space is seeded with “cortical glue” agents that bind the astral microtubules tightly to mimic the observation that once spindles rotate one pole to the cortex they remain stuck. Finally, 2000 dynein motors, which can bind cortical microtubules (mimicking dynein at the cortex) but also bind and motor along astral microtubules (mimicking spindle pole dynein), are added to the entire simulation space and allowed to diffuse around freely. Dyneins, represented as red dots, bind microtubules (white lines) and pull the spindle along. If the spindle reaches the cortex with its astral microtubules they are bound by the “cortical glue” (purple dots). Each simulation is allowed to run for 300 simulated seconds (300,000 frames at 1 millisecond per frame).

## ACKNOWLEDGEMENTS

We thank Reto Gassmann, Sander van den Heuvel and Paul Mains for strains. Some strains were provided by the Caenorhabditis Genetics Center, which is funded by NIH Office of Research Infrastructure Programs (P40 OD010440). We thank Francois Nedelec for help utilizing Cytosim.

## AUTHOR CONTRIBUTIONS

E.V., K. M. and M.T.P: design and execution of experiments and analysis of results. D.B.C.: Mathematical modeling. A.S.M and F.J.M: Experimental design, data analysis and writing.

## SUPPLEMENTARY INFORMATION

Cytosim code

Movie 1: Flat poles, short vs long axial ratio, short astral microtubules.

Movie 2: Pointed poles, short vs long axial ratio, short astral microtubules.

Movie 3: Flat poles, short vs long axial ratio, longer astral microtubules.

Movie 4: Pointed poles, short vs long axial ratio, longer astral microtubules.

Movie 5: Flat poles, short vs long axial ratio, longer astral microtubules, reduced drag.

